# Selecting for lysis in a temperate phage population

**DOI:** 10.64898/2026.10.07.757260

**Authors:** Haley Atkins, Catherine Putonti

## Abstract

Temperate phages represent an abundant source of phage diversity, but their capacity for lysogeny limits their therapeutic potential. Here, we asked whether repeated selection for lytic replication could drive a temperate phage toward an obligately lytic lifestyle. We serially propagated a cocktail of three temperate coliphages using a modified Appelmans’ protocol in which phage populations were independently passaged on eight *E. coli* hosts. Whole-genome sequencing revealed substantial changes in population composition and mutation frequencies. The Uetakevirus phage 1362 was undetectable after the first round, whereas the closely related P2-like phages in the cocktail dominated the evolving populations. Mutations repeatedly accumulated in lysis- and lysogeny-associated genes. Longitudinal genomic analysis revealed rapid fixation of mutations in the integrase coding region, while variants in *lysA*, *lysB*, and *holin* increased in frequency over successive rounds of selection. Protein structure modeling localized selected mutations to regions with potential functional significance. These include the DNA-binding domain of integrase and a surface-exposed region of LysA. Together, these results show that sustained selection for lytic replication can drive rapid and repeatable genomic adaptation in temperate phage populations.

**Importance:** Temperate phages could substantially expand the pool of candidates for phage therapy if their ability to enter the lysogenic life cycle can be overcome. While genetic engineering has successfully produced obligately lytic derivatives of temperate phages, experimental evolution has rarely been applied to directly select temperate phage populations towards increased lytic replication. Here we show that repeated selection for lytic replication drives repeatable genomic changes in temperate coliphage populations, including rapid fixation of mutations in the integrase and accumulation of variants in lysis-associated genes. Our study expands the traditional application of the Appelmans’ protocol beyond phage host-range expansion and establishes a framework for directing naturally abundant temperate phages toward increased lytic activity.

## Introduction

The escalating burden of antibiotic resistance has renewed interest in phage therapy as a complementary approach to combating multidrug-resistant bacterial infections (1, 2). Phage therapy has shown promising efficacy in the treatment of bacterial infections across various anatomic sites, clinical presentations, and bacterial species (see reviews (3–5)). Because individual phages often exhibit narrow host ranges, phage therapy selectively targets pathogenic bacteria without disturbing beneficial members of the microbiota, thus reducing the risk of collateral damage often seen with broad-spectrum antibiotics (6). On the other hand, isolating and validating phages for therapeutic use can be laborious and time-consuming (7).

Experimental evolution has been harnessed to broaden the host range of putative therapeutic phages. The Appelmans’ protocol is an *in vitro* evolution method in which phage cocktails are serially propagated on a collection of bacterial isolates until the evolved phage population can lyse previously resistant strains (8). This protocol has successfully expanded the host range of phages targeting a variety of bacterial pathogens, including *Pseudomonas aeruginosa* (8)*, Enterococcus faecium* (9)*, Acinetobacter baumannii* (10), and *Klebsiella pneumoniae* (11). To date, the Appelmans’ protocol has been applied almost exclusively to virulent (strictly lytic) phages to expand their therapeutic host range, while its application to temperate phages remains largely unexplored.

Lytic phages are preferred for therapeutic use because they do not integrate their genetic material into the bacterial genome, minimizing risks associated with lysogenic conversion, and increasing the likelihood that the target pathogen will be effectively killed. Temperate phages are generally avoided because of their ability to mediate horizontal gene transfer, potentially facilitating the spread of antibiotic resistance and virulence genes (12). However, temperate phages are abundant in nature; nearly 50% of sequenced bacteria to date are lysogens (13). Instead of labor-intensive “phage hunting” for obligately lytic phages, sequencing allows us to rapidly identify temperate phages. Recent advances in synthetic biology have made it possible to engineer temperate phages using a wide range of molecular tools (see reviews (14, 15)). Additionally, genes required for the establishment and maintenance of lysogeny can be modified or deleted (16, 17). As a result, temperate phages can be converted into obligately lytic phages with therapeutic potential.

An alternative approach is to experimentally evolve temperate phages under conditions that favor repeated lytic replication. While experimental evolution has been used to broaden phage host range, its potential to reshape phage lifestyle has received comparatively little attention. Early experimental work identified mutations in the lambda phage CI repressor, which represses lytic development, following selection for lytic activity (18). To investigate whether sustained selection could shift temperate phages toward lytic replication, we modified the traditional Appelmans’ protocol to select for lytic efficacy. Rather than pooling lysates after each round of propagation, lysates were returned only to the host strain from which they were recovered. Here we used a phage cocktail consisting of three temperate phages – 1091 and 1162, which are both P2-like (*Peduovirus*) phages, and 1362, a *Uetakevirus*. Phages 1091 and 1162 were chosen because of their genomic similarity; the two genomes only differ at 46 nucleotides. The work of Burrowes *et al*. (8) and Sáez Moreno *et al*. (19) demonstrated that high sequence similarity between input phages increases the likelihood of homologous recombination. Phage populations were sampled throughout the experiment and analyzed by whole-genome sequencing to identify genetic changes associated with increased lytic efficacy. Mutational changes were tracked over time and exploratory protein structural analysis was performed. By leveraging the abundance and accessibility of temperate phages, this approach provides a workflow for expanding the pool of candidate phages for therapeutic development.

## Results and Discussion

### Selecting for lytic activity

In our previous work with the three temperate phages included in our cocktail (20), we determined that each exhibits a distinct host range (**Table 1**). Only *E. coli* strains lacking detectable prophages were included in the evolution experiment to eliminate native prophage induction, as supported by prior findings (21). The laboratory strain, *E. coli* C, was also included as it was permissive to all three phages prior to the experiment. The lytic activity of the cocktail was assessed every five rounds via spot testing on *E. coli* C (**Table 1**).

**Table 1.**
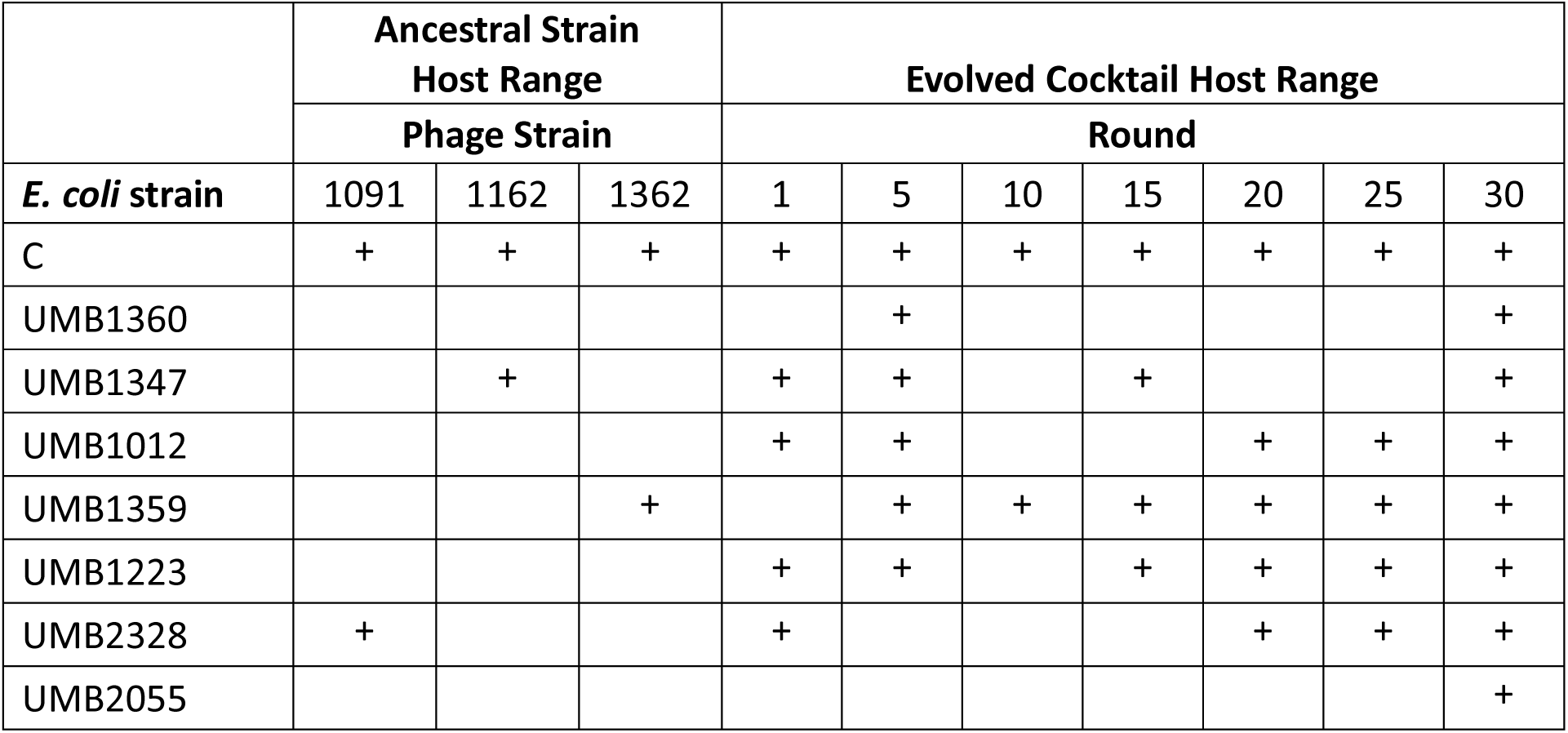
Phage host range for ancestral phages and cocktails evolved on specific bacterial hosts. “+” indicates visible complete lysis (formation of clear plaques); no entry indicates no visible lysis. The “Evolved Cocktail Host Range” corresponds to the lysate collected after the listed number of rounds of selection when tested on the bacterial strain that it had evolved on.

|  | Ancestral Strain Host Range |  |  | Evolved Cocktail Host Range |  |  |  |  |  |  |
| --- | --- | --- | --- | --- | --- | --- | --- | --- | --- | --- |
|  | Phage Strain |  |  | Round |  |  |  |  |  |  |
| *E. coli* strain | 1091 | 1162 | 1362 | 1 | 5 | 10 | 15 | 20 | 25 | 30 |
| C | + | + | + | + | + | + | + | + | + | + |
| UMB1360 |  |  |  |  | + |  |  |  |  | + |
| UMB1347 |  | + |  | + | + |  | + |  |  | + |
| UMB1012 |  |  |  | + | + |  |  | + | + | + |
| UMB1359 |  |  | + |  | + | + | + | + | + | + |
| UMB1223 |  |  |  | + | + |  | + | + | + | + |
| UMB2328 | + |  |  | + |  |  |  | + | + | + |
| UMB2055 |  |  |  |  |  |  |  |  |  | + |

We selected four *E. coli* strains with no evidence of lysis by any of the three ancestral phage strains. By Round 5, the phage cocktail could completely lyse three of these strains (*E. coli* UMB1360, *E. coli* UMB1012, and *E. coli* UMB1223), suggesting rapid adaptation. Lysis via the cocktail was observed for the fourth strain, *E. coli* UMB2055, following Round 30. As the cocktail persisted throughout the course of the experiment, this suggests that lysis occurred throughout; however, it was not captured via our spot tests consistently until Round 30. The intermittent lysis patterns seen for some bacterial host strains further suggest ongoing shifts in phage population dynamics, possibly driven by selection pressures within each round of the protocol. For instance, *E. coli* UMB1359 was initially susceptible to phage 1362, although lysis was not observed by the cocktail after Round 1. Nevertheless, it showed complete lysis in all subsequent spot tests (**Table 1**).

### Genomic Analysis of Evolved Phage Cocktails

To investigate genetic changes associated with enhanced lytic activity, phage populations following Rounds 1, 5, 10, 15, 20, 25, and 30 were sequenced. A total of 46 single nucleotide polymorphisms (SNPs), across multiple coding regions, differentiate the ancestral sequences of phage 1091 and phage 1162. Given the sequence similarity of the two P2-like phages, we selected 1162 as a representative for this population. From the earliest time point (Round 1), no detectable reads aligned to the *Uetakevirus* phage 1362 ancestral genome sequence. This was consistent across all hosts and suggests either integration (although not expected for the *E. coli* C host) or more likely rapid displacement or extinction of 1362 from the cocktail population.

Although this outcome was unexpected given the initial equal titers of phage in the initial cocktail, the rapid loss of phage 1362 is consistent with strong selective pressure between two closely related phages. Similar loss of individual phages has been reported during Appelmans’ evolution experiments, where differences in phage fitness lead to changes in cocktail composition, including the loss of individual phages through competitive outgrowth by other members of the cocktail (8, 10). Phage 1362 may have exhibited slower replication dynamics, a longer latent period, or delayed adaptation to the experimental conditions, allowing phages 1162 and 1091 to dominate the population before phage 1362 could establish itself (10). Thus, the observed extinction of phage 1362 likely reflects differences in competitive fitness during serial propagation.

Overall, each of the eight phage cocktails could lyse its respective bacterial strain by Round 30, despite intermittent observed plaquing (**Table 1**). This suggests that the phages were able to persist through periods of reduced lytic activity, which may reflect periods of partial lysis or low-level activity not captured in our sequencing data, as only instances of complete lysis were investigated. In several cases, phages that produced visible, clear spots on overlays could not be successfully extracted or sequenced from the corresponding lysates, suggesting some limitations in DNA yield, extraction efficiency, or sample degradation. For example, sequencing of the harvests from the cocktails propagated on *E. coli* UMB1360 (Round 25), *E. coli* UMB1359 (Rounds 10 and 30), *E. coli* UMB1347 (Rounds 25 and 30), and *E. coli* UMB1223 (Rounds 25 and 30) did not result in any detectable phage reads, indicative of spontaneous loss of phage populations from the lysates.

### Mutations after One Round of Selection

First, we considered the mutations observed after 1 round (8 hours of culture) of the experiment. For the sake of the discussion here, we will refer to these metrics of variance to describe the mutation frequencies: low variability as ∼0-30%; intermediate variability as ∼40- 60%; high variability as ∼70-100%. The sequence variation for phage 1162 propagated through the different hosts is shown in **Figure 1**. This analysis revealed substantial heterogeneity in mutation frequencies between the hosts, suggesting early adaptation events or pre-existing diversity in the phage population. This variation exceeded the differences observed between the genome sequences of the ancestral phages 1091 and 1162. Notably, many mutations were already present at moderate to high frequencies after Round 1 of the experiment (dark colors, **Figure 1**).

**Figure 1.**
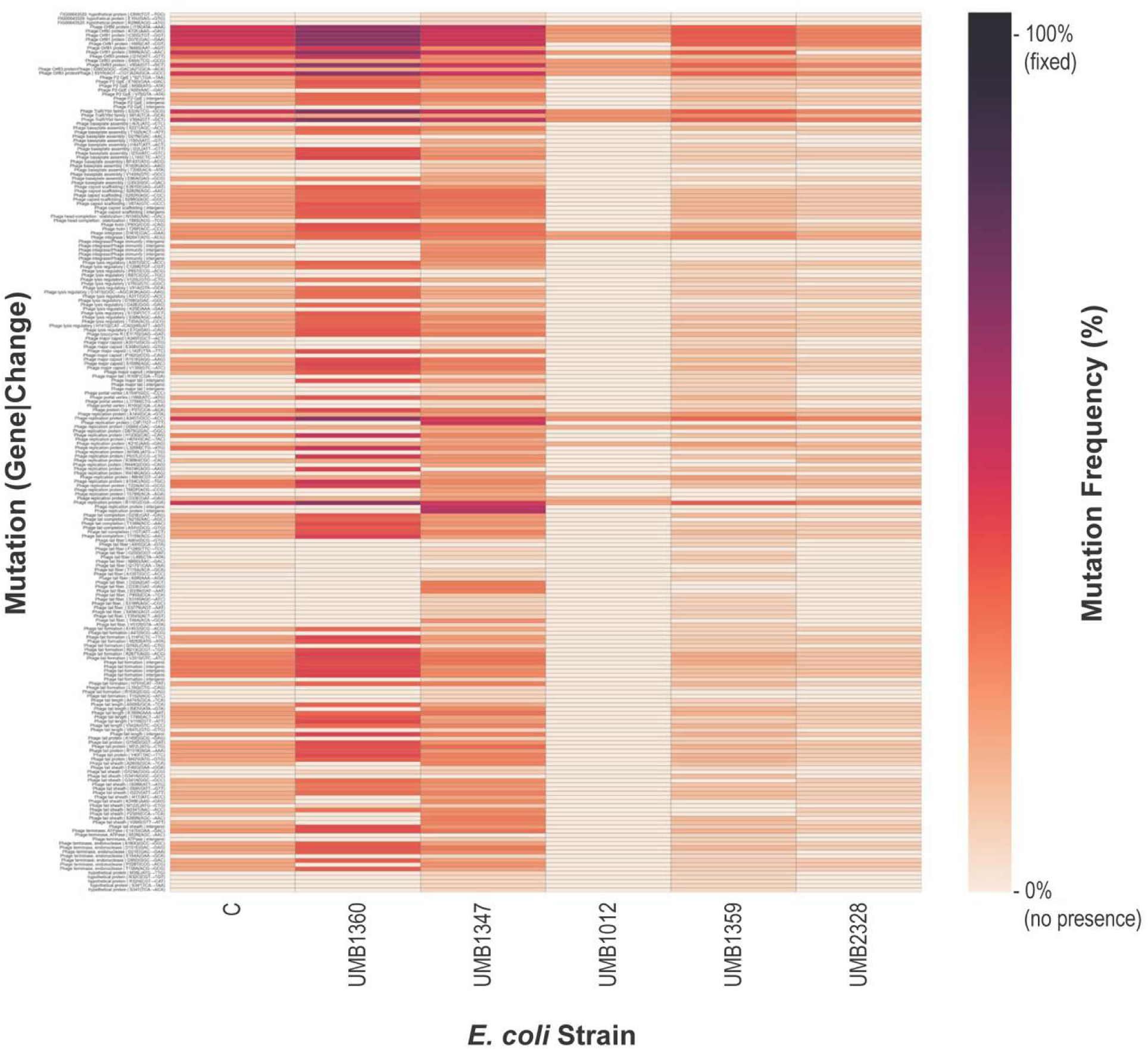
Mutations of phage population across hosts after Round 1. Each bar represents a nonsynonymous mutation or mutation within an intergenic region identified in the phage genome for each host. Along the Y axis are the mutated gene/ region, and along the X axis are the different hosts. Mutations are plotted by genomic position and gene, with variant frequencies indicated by color. Light orange is 0% frequency within the population and black is 100% frequency within the population (fixed).

The high (40-50%) frequency mutations observed for the cocktails propagated through *E. coli* C, *E. coli* UMB1360, and *E. coli* UMB1347 after Round 1, could reflect variation in the ancestral phage stock or *de novo* mutations that increased fitness within the new host. For instance, after Round 1 in *E. coli* UMB1347, the phage sequence includes nonsynonymous substitutions affecting baseplate assembly, tail fiber proteins, and replication-associated genes. These mutations were found exclusively or at elevated frequencies in phages passaged through this *E. coli* host. Several of these same mutations are observed in the phage populations recovered from *E. coli* UMB1359 and *E. coli* UMB2328, albeit typically at a significantly lower frequency within the population. However, we do not see such variation in the cocktail passaged through *E. coli* UMB1012. One could speculate that most phage variants were deleterious within the *E. coli* UMB1012 host.

The presence of high-frequency mutations after just one round of selection raises important considerations about their origin. Although rapid selection can drive such changes, the shared presence and similar frequencies of many mutations across phages passaged through different hosts suggest that some polymorphisms were likely present in the initial phage population. This pre-existing variation, rather than de novo mutation alone, could explain the immediate emergence of common mutations across diverse hosts as observed in other studies (22). Additionally, a key feature of the experimental design was the pooling of phage from multiple independent wells at each transfer step. This pooling increased the mutational input by expanding the opportunity for different variants to arise and persist across replicates. Because each row consisted of a 10-fold (10^0^ – 10 ^−9^) dilution series, applied consistently each round, the initial phage population size, and thus the strength of genetic drift versus selection, varied substantially between wells. In wells with the highest dilutions, bottlenecks were likely severe, amplifying the effects of drift, whereas lower dilutions maintained larger populations more amenable to selection. These dynamics, coupled with the pooling approach, may have facilitated both the retention of rare variants and the rapid emergence of beneficial ones.

These early dynamics underscore the importance of tracking mutation trajectories over time, while also highlighting the limitations of this approach: mutations that arise on the same genetic background will shift in frequency together, making it difficult to distinguish genuinely adaptive variants from neutral hitchhikers. Although recombination can help separate linked mutations, its effectiveness depends on recombination rate, mechanism, and the physical distance between loci on the genome.

### Temporal Mutation Trajectories

**Figure 2** shows the mutations observed over the course of the selection experiment for the phage cocktail propagated through the host *E. coli* C, again with darker colors signifying their predominance in the population. A significant number of variants in tail-related coding regions reached fixation early in the selection experiment, suggesting strong pressure on structural components critical to *E. coli* C host recognition and infection. A full list of fixed mutations is included in **Table 2**. Also worth noting, mutations within the small tail structural protein *GpU* were observed, but they either did not fix within the 30 rounds of selection performed or were lost by Round 30, suggesting they may have been deleterious, neutral, or beneficial but outcompeted due to clonal interference or linkage to less fit genetic backgrounds.

**Figure 2.**
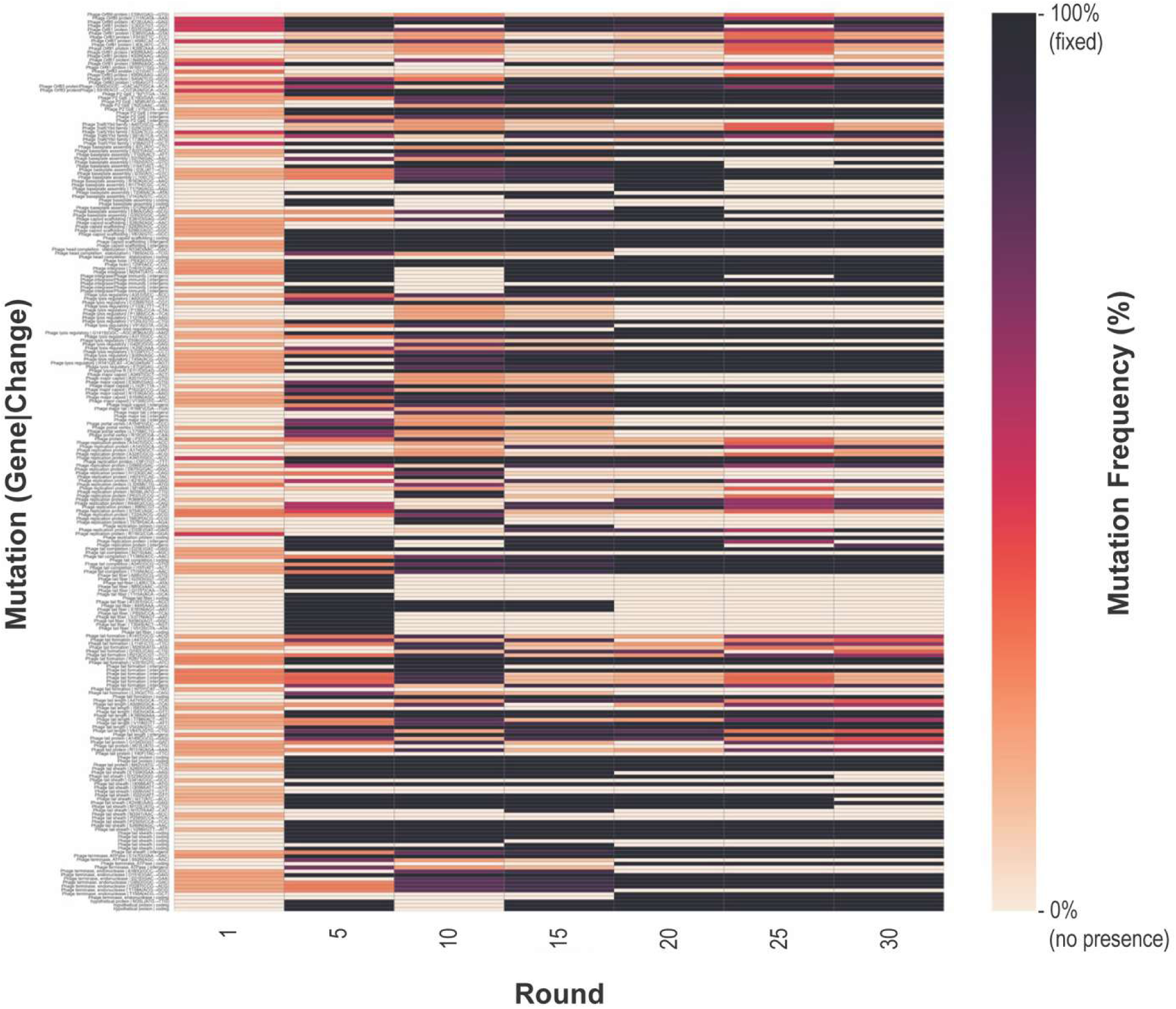
Mutations across phage populations evolved on host *E. coli* C. each bar represents a nonsynonymous mutation or mutation within an intergenic region identified in the phage genome. Along the Y axis are the mutated gene/ region, and along the X axis are the days. Mutations are plotted by genomic position and gene, with variant frequencies indicated by color. Light orange is 0% frequency within the population and black is 100% frequency within the population (fixed).

**Table 2.** Mutations that fixed in tail-related coding regions of the cocktail propagated through the *E. coli* C host.

| Predicted Function | Round |  |  |  |
| --- | --- | --- | --- | --- |
|  | 5 | 10 | 15 | 20 |
| Tail Formation protein *GpD* | R267T, V201I |  |  |  |
| Tail Fiber Assembly<br>- *GpG* | A66V, G25D, L49I, N89D, T115A |  |  |  |
| Tail Completion<br>- *GpR*<br>- *GpS* |  |  |  | T138N, A54V, T115N |
| Baseplate Assembly<br>- *GpJ*<br>- *GpV* |  |  | T102I<br>I150V | V142A |
| Tail Length tape-measure |  | V542A, K186N | I583V |  |
| Tail Proteins<br>- *GpX* | M42V |  |  |  |

For the phage cocktails passaged through the clinical *E. coli* hosts, we were unable to recover the viral genomic sequence from all seven time points. This is not surprising given the intermittent plaquing observed (**Table 1**). For the phage passaged through *E. coli* UMB1347 (**Figure S1**), the P2-like phage was recovered via sequencing after Rounds 1, 10, and 15. Just as was observed for the cocktail passaged through *E. coli* C, we identified mutations accumulating and fixing within tail-associated genes (**Table S1**). Furthermore, we observed variants in the population present in early rounds that were swept from the population. These likely represent transient variants, either under weak selection or outcompeted by other mutants later in the experiment. For the phage cocktail propagated on *E. coli* UMB1012, we were able to assess variants emerging within the population, comparing the genomic data from Round 1 and Round 10 (**Figure S2**). A notable trend was the swift rise in mutations across numerous tail-associated genes, including tail formation, tail completion, tail sheath, and tail length; multiple variants absent at Round 1 were fixed by Round 10 (**Table S2**). For the phage population passaged through host *E. coli* UMB2328, only Round 1 and Round 30 samples were successfully sequenced (**Figure S3**). Several key tail fiber mutations that were fixed in the Round 30 data were absent at Round 1 (**Table S3**). Additionally, nonsynonymous mutations in the tail sheath, tail length, and baseplate genes were observed in the Round 30 genomic data (**Table S3**).

Furthermore, variants within these genes present at low frequencies at Round 1 were absent by Round 30, suggesting transient, non-beneficial mutations or neutral hitchhikers that were outcompeted by more adaptive lineages as selection progressed.

### Adaptive Changes in Lysis-Associated Genes

Regardless of the host for which the cocktail was propagated, mutations within genes associated with lysis were routinely observed. This supports our claim that serial propagation under the modified Appelmans’ protocol selects for increased lytic behavior independent of host-specific adaptation. To better understand the genetic changes driving enhanced lytic activity during experimental evolution of phage 1162, we next focused on genes directly involved in regulating the phage lifecycle—*lysA*, *lysB,* holin, and integrase. Genes *lysA* and *lysB* are well characterized within the P2 virus lysis module (23). The *lysA* gene product functions as an antiholin, regulating the timing of holin activity to ensure lysis occurs at the optimal moment. *lysB* and *lysC* are homologous to the Lambda phage spanin components *Rz* and *Rz1*, which function to fuse the inner and outer membrane during the final stage of lysis (23).

Our analysis focused on phage populations propagated on hosts *E. coli* C and *E. coli* UMB2328, as these are the two phage cocktails that had timepoint data by Round 30. Because *lysA*, *lysB*, and holin play key roles in host-cell lysis and phage release, while integrase enables the establishment of lysogeny through site-specific integration of the phage genome, sequence variation in these genes may influence phage–host interactions, lysis efficiency and timing, and lysogenic behavior (24, 25). **Figure 3** shows the temporal progression of nonsynonymous mutations within and regions adjacent to the *lysA*, *lysB*, holin, and integrase coding regions for the cocktail passaged through *E. coli* C (panel A) and *E. coli* UMB2328 (panel B). A clear pattern emerged: mutations in the integrase gene became fixed early, while mutations in *lysA* and *lysB* accumulated progressively, suggesting strong directional selection for enhanced lytic behavior and concurrent loss of lysogeny-related functions.

**Figure 3.**
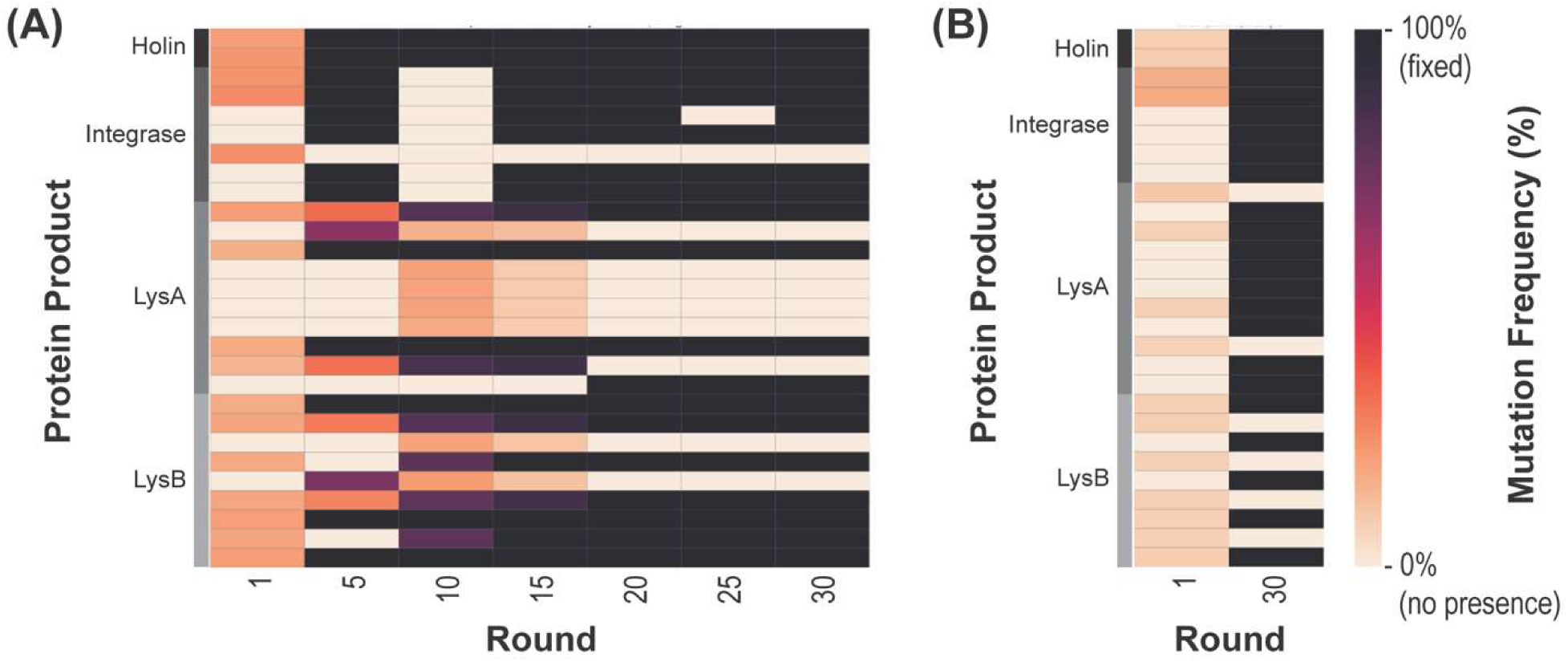
Nonsynonymous mutations and regions identified in (or adjacent to) *lysA*, *lysB*, *holin*, and *integrase* genes in phages evolved on *E. coli* C (A) or *E. coli* UMB2328 (B). Mutations are plotted by amino acid substitution within each gene. The Y axis includes a gray scale bar indicating if the coding region containing the mutation, with variant frequencies indicated by color. Light orange is 0% frequency within the population and black is 100% frequency within the population (fixed).

In particular, two integrase missense mutations—D161E (GAC→GAA) and M264T (ATG→ACG)—rose to fixation rapidly in the phage populations on both *E. coli* hosts. Though absent on Round 1, they were fixed by Round 5 in the *E. coli* C phage population, with only a brief absence observed at Day 10 before reappearing at 100% frequency by Round 15. This rapid fixation indicates a strong early selective sweep likely inactivating or impairing integrase function, favoring a lytic strategy. The early timing and parallel fixation of these mutations suggest that loss of lysogeny conferred an immediate advantage under the experimental conditions, as observed in other studies (26). These two mutations were also fixed at Round 30 in the cocktail passaged through *E. coli* UMB2328. Because no other data points are available for this host, we are unable to determine if these mutations fixed early in the selection experiment.

While D161E represents a conservative substitution, M264T represents a more dramatic shift within the DNA-binding domain, which could significantly alter integrase recognition and binding to target DNA (**Figure 4**). While direct experimental data on these specific mutations are lacking, prior work on phiC31 integrase demonstrates that mutations in conserved regions— particularly those within or near the DNA-binding domain—often abolish recombination activity by impairing DNA binding or strand cleavage (27). These findings support the hypothesis that the observed integrase mutations compromise its ability to mediate prophage integration, favoring a lytic replication strategy.

**Figure 4.**
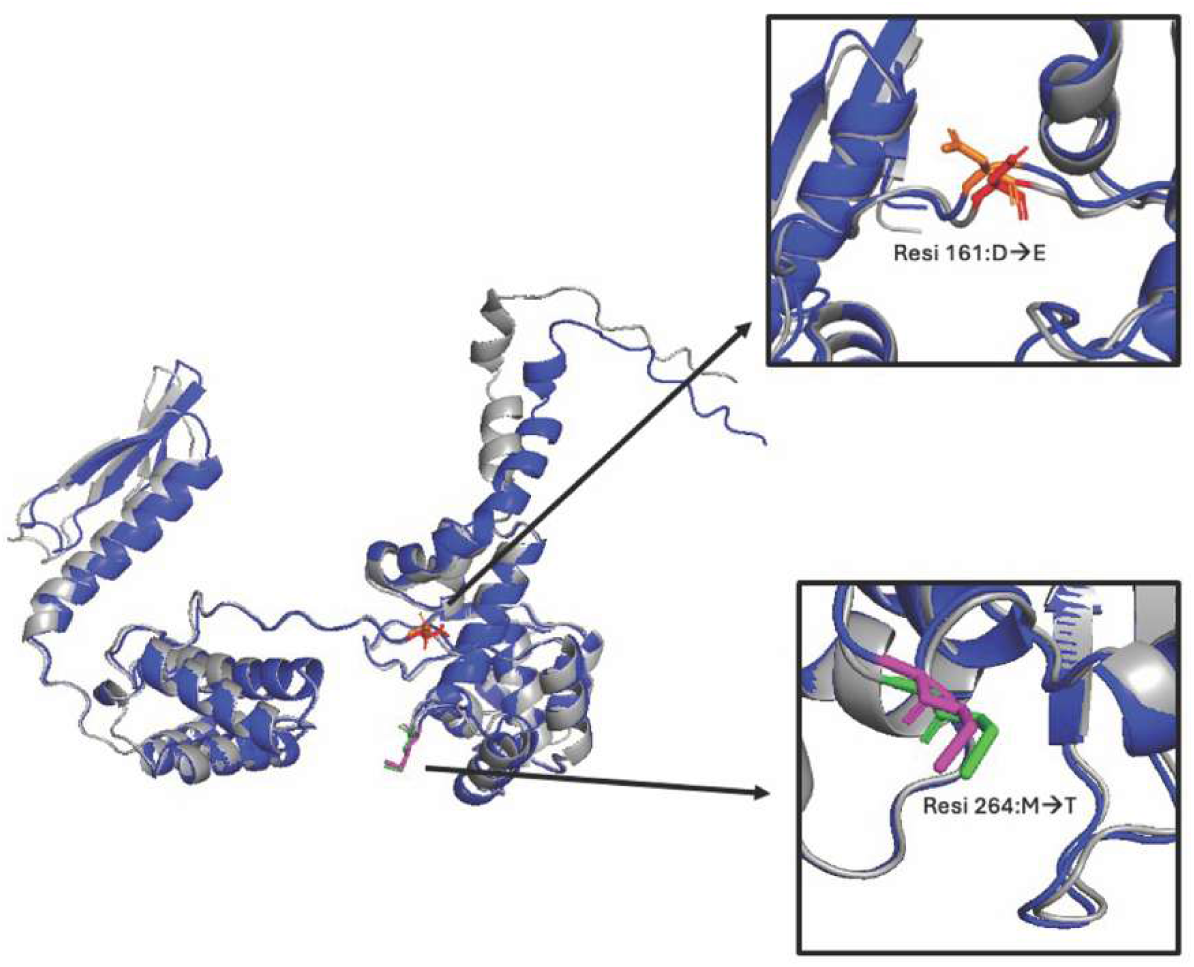
Three-dimensional structural comparison of the wild-type and mutant integrase proteins highlighting two nonsynonymous mutations for the phage cocktail on *E. coli* C or *E. coli* UMB2328. WT integrase protein (grey) aligned with the mutant integrase (blue) using PyMOL (RMSD=0.625Å). Each mutation site is depicted using stick representation, with the WT distinctly colored from the mutant. (D161E): Wild-type aspartic acid (red) is substituted with glutamic acid (orange) at residue 161; (M264T): Wild-type methionine (green) is substituted with threonine (magenta) at residue 264.

In *lysA*, phage 1162 also displayed strong signals of selection. Mutations P85T (CCG→ACG), R87C (CGC→TGC), P138S (CCA→TCA), P138L (CCA→CTA), and V120L (GTG→CTG) were all present at Round 1, albeit as minor variants. However, they were fixed in the population at Round 30 for *E. coli* UMB2328, our only other time point for this host (**Figure 3B)**. V120L fixed earlier (Round 5) in the cocktail passaged through *E. coli* C (**Figure 3A**). As **Figure 5** shows, these mutations cluster within a surface-exposed region of the protein, suggesting these residues may participate in substrate binding or interaction with host cell components.

**Figure 5.**
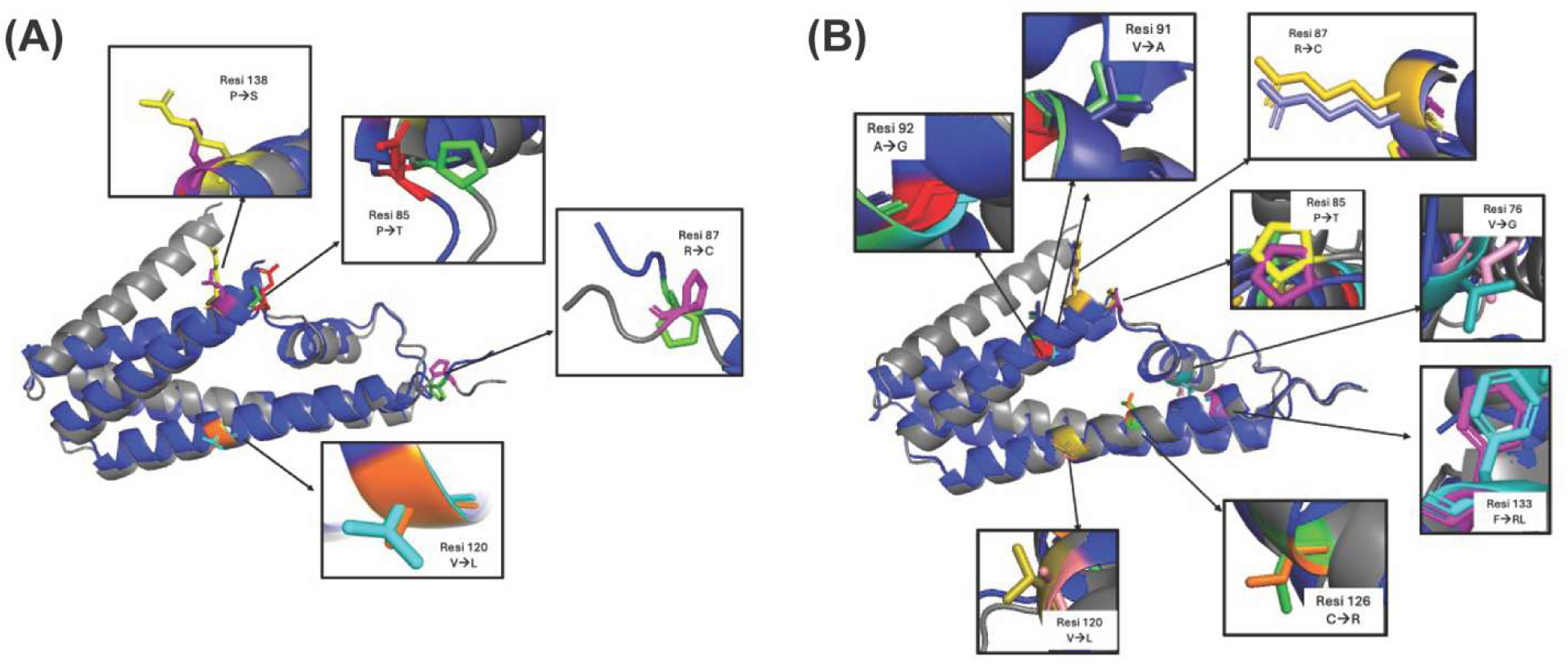
Three-dimensional structural comparison of the wild-type and mutant LysA proteins highlighting nonsynonymous mutations for the phage cocktail propagated through **(A)** *E. coli* C and **(B)** *E. coli* UMB2328. WT LysA protein (grey) aligned with the mutant LysA (blue) using PyMOL (RMSD=1.156Å). Each mutation site is depicted using stick representation, with the WT distinctly colored from the mutant. In panel A, (P138S): Wild-type proline (yellow) is substituted with serine (purple) at residue 138; (P85T): Wild-type proline (green) is substituted with threonine (red) at residue 85; (R87C): Wild-type arginine (magenta) is substituted with cysteine (green) at residue 87; and (V120L): Wild-type valine (orange) is substituted with leucine (cyan) at residue 120. In panel B, (A92G): Wild-type alanine (blue) is substituted with glycine (red) at residue 92; (C126R): Wild-type cysteine (green) is replaced by arginine (orange) at residue 126; (F133L): Wild-type phenylalanine (cyan) is mutated to leucine (magenta) at residue 133; (P85T): Proline (yellow) is replaced by threonine (purple) at residue 85; (R87C): Arginine (slate) is substituted with cysteine (gold) at residue 87; (V120L): Valine (olive) is replaced with leucine (salmon) at residue 120; (V76G): Valine (teal) is mutated to glycine (pink) at residue 76; and (V91A): Valine (lime) is substituted by alanine (deep blue) at residue 91.

Mutations that fixed in both selection experiments, such as P85T and R87C, are located adjacently on the protein surface, highlighting a potential hotspot for functional adaptation. In contrast, the additional substitutions seen in the phage propagated through *E. coli* UMB2328 (**Figure 5B**) extend this cluster and introduce further chemical diversity to the protein surface. These findings suggest that LysA’s activity possibly affects substrate binding, enzymatic function, or host-specificity. This is consistent with studies showing that phage lytic enzymes often undergo rapid adaptation in response to host resistance (28).

Mutations in the *lysB* gene displayed distinct host-specific patterns. In the phage passed through *E. coli* C, several mutations became fixed at different time points: G141A and S38N by Round 5, T45A and G42E by Round 15, S135P and A31T by Round 20. Furthermore, an early stop codon was detected in our Round 10 sequencing data. This truncation likely disrupts the full spanin domain, potentially impairing the ability to coordinate with holin-endolysin systems during lysis. In contrast, variants from the phage cocktail passaged through host *E. coli* UMB2328 exhibited a different set of mutations that were fixed by Round 30: K25E, G42E, and D106G. Based upon their location within the protein structure (**Figure S4**), we hypothesize that D106G likely alters flexibility and charge distribution within the C-terminal region of the spanin domain, while K25E replaces a basic lysine with an acidic glutamic acid near the N-terminus, possibly tuning membrane interactions through surface charge. Furthermore, many of the mutations that swept in the phage population passaged through *E. coli* C were also present at low frequency in the phage population passaged through host *E. coli* UMB2328. This pattern suggests that while both populations may have shared a common genetic background, the *lysB* mutations in the phage cocktail passaged through *E. coli* UMB2328 appear to originate from a different background, indicating possible recombination events introducing new combinations of mutations.

Mutations in the holin—**P93Q** and **T29P**—were observed to fix in the phage propagated through both host *E. coli* C (by Round 5) and host *E. coli* UMB2328 (by Round 30), suggesting selection across diverse host environments. The parallel fixation of these mutations across hosts suggests a conserved adaptive strategy; however, incomplete time points prevent assessment of whether fixation occurred synchronously. Structural modeling in PyMOL showed that the mutated holin protein aligns perfectly with the reference structure (figure not shown). This suggests that **P93Q** and **T29P** do not significantly disrupt the overall conformation of the protein. Instead, these mutations may influence holin function through subtle mechanisms. Prior work has found that different holin alleles confer different lysis times in λ phage (29).

## Conclusions

Many of the mutations observed are localized to genes involved in host recognition, DNA replication, or lytic cycle regulation, consistent with known targets or selection during experimental evolution and phage engineering (30). Collectively, these findings reveal a strong pattern of adaptive evolution within lysis-associated genes in phage 1162, marked by early fixation of integrase mutations and the rise of multiple missense mutations in *lysA* and *lysB*. However, these mutations occurred on a shared genetic background that likely included additional beneficial mutations in other genes. While this background was advantageous during repeated passage through high-density bacterial cultures, where rapid host clearance confers a competitive edge, it remains unclear whether the lysis-related mutations themselves were directly beneficial or merely hitchhiked with other adaptive changes. The recurrence of these mutations across timepoints and hosts, despite differences in host background, suggests homologous recombination facilitated the spread of this beneficial genetic background. At the same time, the presence of unique mutations in later rounds or on specific hosts (e.g., host *E. coli* UMB2328) reflects ongoing diversification and fine-tuning, hallmarks of adaptive phage evolution under dynamic selective regimes (31).

Experimental evolution of phage populations across multiple *E. coli* hosts revealed evidence of adaptive mutation, particularly within genes encoding structural and lysis-related proteins. Across phages propagated in hosts *E. coli* C, *E. coli* UMB1347, *E. coli* UMB1012, and *E. coli* UMB2328, tail-associated genes exhibited strong signs of selection, with many mutations rapidly reaching fixation, suggesting their critical role in overcoming host-specific defenses.

Parallel evolution in lysis-regulating genes indicated a shift toward lytic behavior, particularly with early fixation of disruptive integrase mutations. In silico protein modeling of these variants suggested potential impacts on enzymatic function, membrane interactions, and host recognition. However, these interpretations remain speculative without direct experimental validation.

Together, these results suggest that phage evolution follows constrained yet repeatable adaptive pathways despite differences in the host environment. Such parallel evolution is consistent with previous phage evolution experiments, indicating that adaptation may be strongly constrained by the accessibility and fitness effects of a limited set of beneficial mutations (32, 33). At the same time, the coexistence of shared genetic backgrounds with host-specific and late-arising mutations highlights the balance between convergence and diversification during phage adaptation. While the fitness effects of individual mutations remain unresolved, the observed patterns are consistent with selection acting on linked suites of traits rather than a single trait in isolation. Overall, this work underscores how experimental evolution can illustrate the structure of phage adaptive landscapes and the evolutionary processes shaping phage-host interactions.

## Methods

### Bacterial and Phage Strains

The *E. coli* C laboratory strain and the previously described 66 *E. coli* strains of Garretto et al. (34) were used to assess host range for the three phages. These 66 *E. coli* strains are urinary isolates, obtained from females with or without lower urinary tract symptoms as part of prior IRB-approved and published studies (35–42). The three phages used in this experiment, phages 1362, 1162, and 1091 are temperate phages, previously isolated via spontaneous induction on the *E. coli* C host (20).

To assess lysis, phages were spotted (10 µL) onto lawns of *E. coli* C. Each lawn consisted of 500 µL of overnight *E. coli* culture, grown at 37°C with shaking, and 3 mL of soft LB agar (0.7% agar) mixed and spread onto a LB agar plate. The spot plates were incubated overnight at 37°C. Evidence of clear plaques was then noted as full lysis.

### Modified Appelmans’ Protocol

To drive evolution, we modified the original Appelmans’ protocol developed at the George Eliava Institute of Bacteriophage, Microbiology, and Virology (8). The protocol is represented schematically in **Figure 6**. Phages 1362, 1162, and 1091 were combined 1:1:1 as an input cocktail. Bacterial strains of *E. coli* were grown overnight in 3 ml LB broth at 37°C with shaking. Using a 96-well microtiter plate, 100 µL of serial 10-fold dilutions (10^0^ to 10^−9^) of the cocktail were added to 100 µL of LB broth and 10 µL of bacterial overnight culture of a single strain plus 100 µL 0.7% saline solution. Eight *E. coli* strains (*E. coli* C, UMB1360, UMB1347, UMB1012, UMB1359, UMB1223, UMB2328, and UMB2055) were used as hosts. After an 8-hour incubation at 37°C on a shaking platform at 200 rpm, the plates were visually inspected, and lysis was documented. Each row, the cocktail dilution for one of the eight *E. coli* hosts, was pooled together and filtered by vortex mixing 1:100 CHCl_3_, followed by centrifugation (15,000 x g for 15 min). This lysate became the round one cocktail and was used to initiate the next round of the protocol using the same bacterial strains. This was repeated 30 times. Every fifth transfer, the lysate was plated on *E. coli* C by spot assays as described above to test for lytic activity.

**Figure 6.**
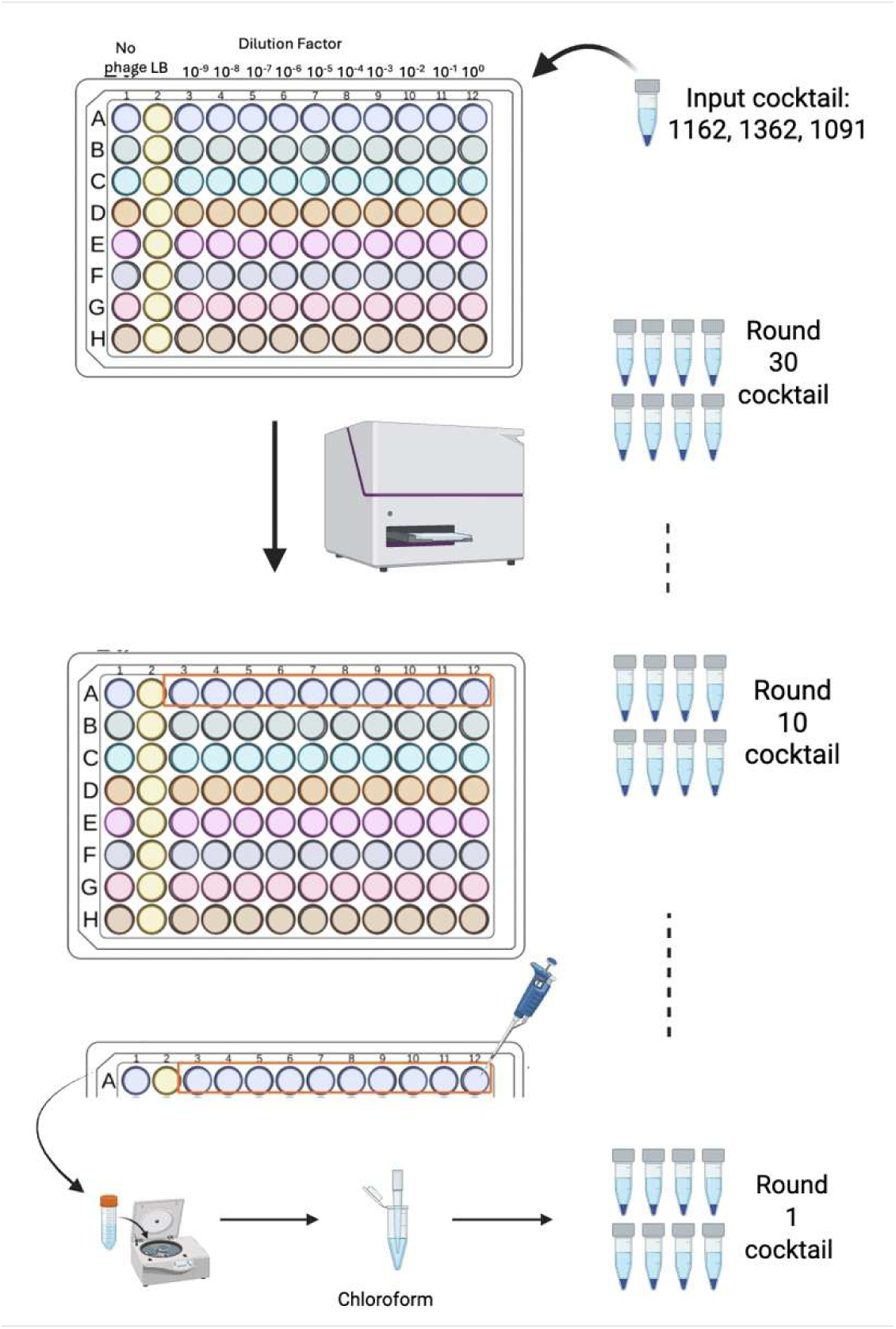
Schematic of the modified Appelmans’ protocol used to drive phage evolution. Three phages (1362, 1162, and 1091) were mixed to create an input cocktail. Eight *E. coli* strains (*E. coli* C, UMB1360, UMB1347, UMB1012, UMB1359, UMB1223, UMB2328, and UMB2055) are represented as A-H. In each round, serial 10-fold dilutions (10⁰ to 10⁻⁹) of the cocktail were added to LB broth, bacterial culture, and saline. Plates were incubated for 8 hours at 37°C, and wells showing lysis were pooled, treated with chloroform, centrifuged, and filtered to produce a new cocktail. This process was repeated for 30 rounds. (Image generated using BioRender.)

### Phage Amplification

To increase phage titers and improve DNA yield, phage amplification was performed using *E. coli* C. 2 mL of LB was inoculated with an *E. coli* C colony from a plate, incubated overnight at 37°C with shaking. 4 mL of LB was added to the overnight culture and incubated for 30 minutes at 37°C with shaking. 100 mL of the Appelmans’ phage lysate was added to the culture and incubated overnight at 37°C with shaking.

### Phage DNA Extraction and Sequencing

The amplified phage was extracted using the Qiagen DNeasy UltraClean Microbial Kit following the manufacturer’s protocol with the following modification. For step one, in a Powerbead tube, 300 mL of Powerbead Solution, 50 mL of Solution SL, and 280 mL phage lysate was added and heated at 70°C for 10 minutes prior to disruption. DNA concentrations were quantified using a Qubit fluorometer. DNA was sent to SeqCenter (Pittsburgh, PA USA) for library preparation and sequencing. Samples were prepared for whole-genome sequencing using an Illumina DNA Prep kit and custom IDT 10 bp unique dual indexes. Sequencing was performed on the Illumina NovaSeq X Plus platform (San Diego, CA, USA) using a 300-cycle flow cell kit, producing a 2×150 bp paired end reads. Demultiplexing, quality control, and adapter trimming was performed with Illumina’s propriety software bcl-convert (v4.2.4) prior to delivery. Raw sequencing reads have been deposited in NCBI under the BioProject Accession No. <u>PRJNA1501873</u>.

### Genome Analyses

Paired-end sequencing reads from phage samples were aligned to each parental reference genome using Bowtie2 (v.2.5.3) (43). Reference genomes were indexed with the bowtie2-build utility. Alignments were performed in default mode. BLAST analysis of unaligned reads confirmed the absence of phage-related sequences.

Whole-genome sequencing data from evolved phage populations (Rounds 1, 5, 10, 15, 20, 25, and 30) were analyzed using the Breseq pipeline (v.0.39.0) to identify mutations relative to the ancestral reference genomes (44). Raw paired-end reads were aligned to the reference genome of the ancestral phage (Phage 1091, 1162, or 1362) using Breseq in polymorphism prediction mode (-p), which is optimized for mixed populations. BV-BRC (v.3.49.1) was used to assemble and annotate the reference genomes (45), and identity was confirmed via NCBI GenBank entries. Mutations were annotated based on their position relative to coding regions and their predicted functional consequences (synonymous, nonsynonymous, intergenic). Breseq outputs were further filtered to focus on lysogeny and lysis genes. Mutated genes of interest (*lysA, lysB,* holin, and integrase) were isolated from the Breseq results and translated into amino acid sequences fasta files via a python script.

### Protein Structure and Domain Analysis

To investigate the potential functional impact of adaptive mutations observed during phage evolution, a combination of protein structure prediction, domain annotation, and molecular visualization was employed. Protein structure prediction was performed using the web version of ESMFold, a deep-learning-based structure predictor (46). Amino acid sequences of phage proteins of interest (integrase, LysA, LysB, and holin) were submitted individually in FASTA format to the ESMFold web server. Predicted models were downloaded in PDB format for downstream visualization and analysis. Molecular visualization and structural interpretation were carried out using PyMol (Schroedinger, LLC). Mutated residues identified by Breseq were mapped onto the predicted protein structures. Conserved domain identification was conducted using the NCBI Conserved Domain Database (CDD) via the CD-Search tool (47). Protein sequences were searched against the CDD v.3.21 database using default parameters, including an E-value of 1.01, low-complexity filtering, composition-based score adjustment, and rescue of borderline hits. Domain boundaries and functional annotations were used to contextualize the location of mutations within known structural or catalytic motifs.

